# A non-destructive approach for measuring rice panicle-level photosynthetic responses using 3D-image reconstruction

**DOI:** 10.1101/2022.06.25.497601

**Authors:** Jaspinder Singh Dharni, Balpreet Kaur Dhatt, Puneet Paul, Tian Gao, Tala Awada, Paul Staswick, Jason Hupp, Hongfeng Yu, Harkamal Walia

## Abstract

Our understanding of the physiological response of rice inflorescence (panicle) to environmental stresses is limited by the challenge of accurately determining panicle photosynthetic parameters and their impact on grain yield. This is primarily due to lack of a suitable gas exchange methodology for panicles, as well as non-destructive methods to accurately determine panicle surface area. To address these challenges, we have developed a custom panicle gas exchange cylinder compatible with the LiCor 6800 Infra-red Gas Analyzer. Accurate surface area measurements were determined with a 3D panicle imaging platform to normalize the panicle-level photosynthetic measurements. We observed differential responses in both panicle and flag leaf for two temperate Japonica rice genotypes (accessions, TEJ-1 and TEJ-2) exposed to heat stress during early grain filling. There was a notable divergence in relative photosynthetic contribution of flag leaf and panicles for the genotype tolerant to heat stress (TEJ-2) compared to the less tolerant accession. The novelty of this approach is that it is non-destructive and more accurately determines panicle area and photosynthetic parameters, enabling researchers to monitor temporal changes in panicle physiology during the reproductive development. The method is useful for panicle-level measurements under diverse environmental stresses, and for evaluating genotypic variation for panicle physiology and architecture in other cereals with compact inflorescences.

## Introduction

Rice (*Oryza sativa*) is a crucial crop for global food security. However, rice production is susceptible to heat stress (HS) (Dhatt *et al*., 2019; Khush, 2005; Khush & Jena, 2009; Muthayya et al., 2014; Prasad et al., 2017; Peng et al., 2004; Moore et al., 2021). Rice reproductive development is considered the most heat sensitive stage (Ali et al., 2019; Arshad et al., 2017; S. V.K. Jagadish et al., 2007; S. V.Krishna Jagadish, 2020; Paul *et al*., 2020a). Even a short duration of heat stress during early grain development affects mature grain size and weight parameters (Chen et al., 2016; Folsom et al., 2014; Kadan et al., 2008; Lisle et al., 2000; Sreenivasulu et al., 2015). During the reproductive stage, rice grain is the primary sink organ whose normal development depends upon the accumulation and utilization of photoassimilates from leaves (Zhang *et al*., 2018; Abdelrahman *et al*., 2020). Recent studies suggest that in addition to being a temporary sink, panicles also contribute to the grain photoassimilate pool and consequently to grain yield (Kong et al., 2016; Tambussi et al., 2021; Vicente et al., 2018a; Chang et al., 2020).

A better understanding of source-sink dynamics in the context of photosynthetic responses and grain filling is needed for predicting how grain yield parameters are affected by temperature (Gao et al., 2021; Lubis et al., 2003; Wang et al., 2020). In absence of further improvement in rice heat resilience, it is estimated that for every 1°C increase in temperature, there will be ∼ 3.2% decline in yield (Zhao *et al*., 2017). From a mechanistic perspective much of that impact could be due to temperature sensitivity of the plant’s photosynthetic capacity and the cellular processes of developing seed. Heat stress impacts photosynthesis in multiple ways, including increasing membrane permeability in leaves, damaging sub-cellular membranes such as thylakoid membranes, thus impeding light harvesting, electron transport rates and ATP generation (Schrader *et al*., 2004; Prasad *et al*., 2008; Djanaguiraman *et al*., 2013; Pokharel *et al*., 2020). Under HS the primary carbon-fixing enzyme, rubisco, is also more active as an oxygenase leading to the production of 2-phosphoglycolate, which is eliminated through the photorespiratory pathway resulting in partial loss of previously fixed carbon (Walker *et al*., 2016; Moore *et al*., 2021). Altogether, the reduced photosynthetic efficiency and increased respiration-photorespiration rates due to HS alter the dynamics between source and sink organs, leading to yield decline (Prasad *et al*.; Ferguson *et al*., 2021).

The capacity of primary source tissue to mobilize photoassimilates and the ability of sink tissue (grain) to accumulate the transported sugars determines the extent of grain filling (Zhang *et al*., 2018; Abdelrahman *et al*., 2020). A significant proportion of the assimilates accumulating in the grains are derived from the upper canopy (Austin et al., 1977; Bidinger et al., 1977.; Inoue et al., 2004). One estimate suggests that the youngest three leaves may contribute over 50% of the assimilates into the rice grain filling pool (Li *et al*., 1998). While foliar tissue is the primary source of photoassimilates, non-foliar tissue such as developing rice panicles are also photosynthetically active and contribute towards photoassimilate accumulation in grains (Tambussi *et al*., 2007; Maydup *et al*., 2012). There is limited evidence supporting the role of green inflorescence tissues as contributor to carbon assimilation (*A*_*gross*_) equivalent to ∼30% of the flag leaf (Grundbacher *et al*., 1963; Imaizumi *et al*., 1990). However, effect of HS on the net photosynthetic contributions by non-foliar organs and the dynamic relationship between foliar and non-foliar organs is not explored in rice. The extent of genotypic variation for these responses is also unknown.

The temporal evaluation of foliar photosynthetic parameters on a per unit area basis can be accomplished non-destructively using well-established protocols. Instrumentation for these experiments is designed for laminar leaf surfaces for which precise surface areas can be determined. However, measurement of non-laminar organs (inflorescence/panicle) with their intricate and complex architectures is challenging (Simkin *et al*., 2020). This issue has been partially addressed recently (Chang *et al*., 2020), where panicle area was computed for evaluating non-foliar photosynthetic parameters. However, this approach limits evaluation of temporal responses due to the destructive sampling of panicles for each measurement. Recent advances in image-based plant phenotyping have enabled the development of a 3D-panicle imaging platform (*PI-Plat*) for high-resolution, temporal assessment of vegetative and inflorescence-related traits in a non-destructive and precise manner (Sandhu et al., 2019; Zhu et al., 2021a; Zhu *et al*., 2021b). Digital traits derived from 3D reconstructed panicles are more sensitive and accurate than results from 2D images (Sandhu et al., 2019). Thus, the non-destructive estimation of panicle size parameters in rice using 3D-imaging platforms can be used to establish surface area normalized panicle photosynthetic assessments.

We combined panicle surface area measurements with a customized gas exchange cylinder that allowed unrestricted enclosure of panicles, thus overcoming a major limitation of shading as reported in other studies (Maydup *et al*., 2012; Sanchez-Bragado *et al*., 2016; Chang *et al*., 2020). Measuring flag leaf and panicle photosynthetic parameters concurrently enabled us to identify relationships between foliar and non-foliar tissue gas exchange rates under control and HS conditions. This novel approach was able to identify changes in source-sink dynamics in response to HS, as well as differential response of two temperate japonica rice accessions that were previously known to differ in their sensitivity to HS during grain development (GSOR Ids: 301110, TEJ-1 and 301195, TEJ-2) (Paul et al., 2020). Our results establish a viable method for more precise temporal evaluation of source-sink relationships during reproductive development, in response to HS, for the study of genetic diversity in photosynthetic strategies among rice accessions. Although we specifically examined HS response, the method should also be useful under other stress conditions as well.

## Materials and Methods

### Plant material and growth conditions

Two temperate japonica rice genotypes, GSOR Ids: 301110 (TEJ-1) and 301195 (TEJ-2), were selected based on their heat stress (HS) response (Paul *et al*., 2020). Mature seeds from the two accessions were dehusked using a Kett TR-130, sterilized with water and bleach (40%, v/v), and rinsed with sterile water. The seeds were germinated in the dark on half-strength Murashige and Skoog media. After five days, germinated seedlings were transplanted to soil in 4-inch square pots, and plants were grown under control greenhouse conditions: 16 h light and 8 h dark at 28±1°C and 23±1°C, respectively. Relative humidity ranged from 55-60% throughout the experiments.

### Heat stress treatments

A set of plants was used to do the PI-Plat imaging and provide photosynthetic measurements using the LI-6800. Plants were grown under control conditions until flowering. Once flowering initiated, the primary panicle was carefully examined. Since the primary panicle is the first one to flower, we focused on measuring its photosynthetic parameters along with flag leaf for experimental accuracy. Upon ∼50% completion of primary panicle flowering, a set of plants was either kept under control conditions (16 h light and 8 h dark at 28±1°C and 23±°C) or moved to greenhouse set-up for a moderate heat stress (HS) treatment (16 h light and 8 h dark at 36±1°C and 32±1°C). We used these plants for primary panicle imaging and photosynthetic measurements at 4 and 10 days after fertilization (DAF) under control and HS conditions (Fig. 4 and S1).

Another set of plants was used for measuring the mature seed yield-related traits. Florets were marked at the time of fertilization to track developing seeds. 1 DAF, plants were kept in either control conditions (16 h light and 8 h dark at 28±1°C and 23±°C) or moved to greenhouse setup for a moderate HS treatment (16 h light and 8 h dark at 36±1°C and 32±1°C). The plants were subjected to HS treatment for either 2 to 4 DAF (HS-I), or 2 to 10 DAF (HS-II). Afterward, plants were moved back to control temperature conditions and harvested at physiological maturity to analyze mature grain yield-related parameters (Fig. S4a).

### Panicle imaging and downstream analysis

#### Image Acquisition

We utilized the Panicle Imaging Platform (PI-Plat) to capture rice panicle images (Gao et al., 2021; Sandhu et al., 2019). Briefly, PI-Plat is comprised of customized wooden chamber (Fig. S1) with a circular wooden board, parallel to the floor, having an aperture at its center. Plants marked for imaging were brought into the chamber, and the primary panicle of the plant was passed through the aperture. A rotary apparatus hosting two Sony α6500 cameras and LED lights (ESDDI PLV-380, 15 watts, 500 LM, 5600 K) rotated 360° around the panicle. With the built-in time-lapse application, each camera took an image per second for one minute. The two cameras generate 120 images for one panicle with a resolution of 6000×4000 pixels. Color checkerboards were placed on the chamber and table to facilitate camera parameters recovery and correspondence detection in paired images (Fuhrmann *et al*., 2014).

#### 3D Point Cloud Reconstruction

Captured panicle images were pre-processed to remove the background. To achieve this, images were first converted from the red, green, and blue (RGB) color space into the hue, saturation, and value (HSV) color space. Then, we implemented color thresholding using the MATLAB application “colorthresholder”. Pixels were removed if their corresponding hue, saturation, and value were not in the range of 0–1, 0–1, and 0.15–1, respectively. Next, the pre-processed images were used for 3D reconstruction. To reconstruct the Panicle’s point cloud, we implemented the Multi-View Environment (MVE) pipeline (Fuhrmann *et al*., 2014). The MVE pipeline detected and matched the image features in the pre-processed images. A parse point cloud was generated based on matched image features. The parameters of cameras, including position and orientation, were also extracted in this process. Afterward, a dense point cloud was generated by calculating the depth information for each pixel in each image using the cameras’ parameters. Finally, floating scale surface reconstruction (FSSR) (Fuhrmann & Goesele, 2014) was implemented to denoise the dense point cloud.

The reconstructed point clouds of the MVE pipeline included all the objects in the scene. We removed uninteresting objects from the original point cloud by implementing color thresholding and connected component labeling to calculate the panicle features in the next section. First, we segmented the panicle’s point cloud cluster by computing the Visible Atmospherically Resistant Index (VARI) (Gitelson *et al*., 2002) for each point in the point cloud. Equation (1) shows the formula of VARI, where R, G, and B mean the corresponding intensity of a point in the RGB color space.

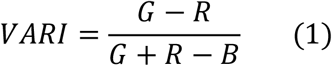

The cluster containing the maximum number of points whose VARI > 0.1 is considered as the panicle. Then, we filtered out uninteresting points in the cluster, for instance, plant labels. A representative image of the final point cloud that includes only the panicle is shown in Fig. 1a.

**Figure 1.**
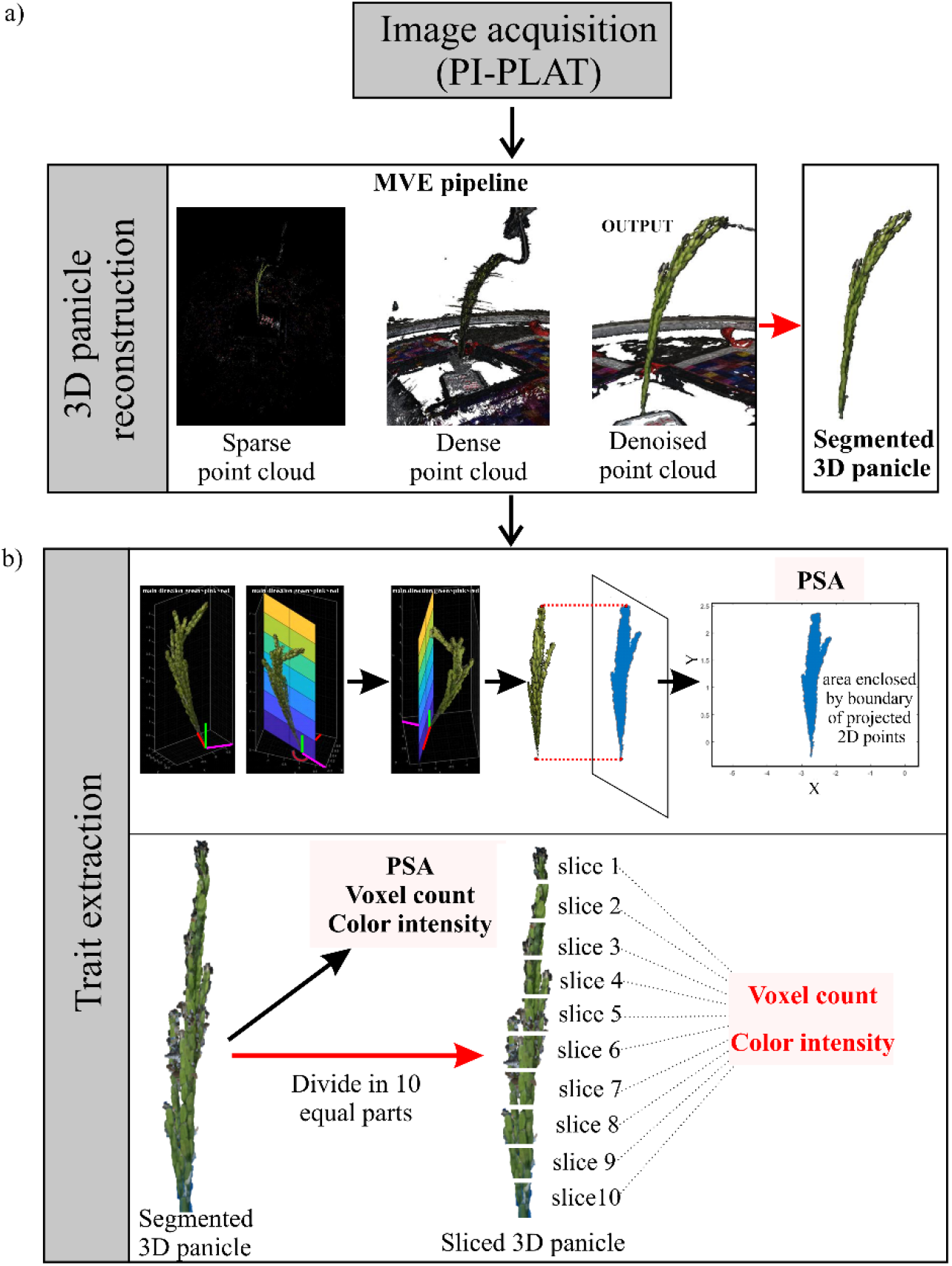
(a) Workflow for reconstruction of 3D panicle from Multiview images using PI-Plat imaging platform. (b) Trait extraction from the reconstructed 3D panicle. The upper panel shows the extracted projected panicle area (PPA) from boundary of projected 2D points. The lower panel shows the traits derived from the segmented 3D panicle and sliced 3D panicle (voxel count and color intensity). Slice 1 corresponds to the top-most slice and slice 10 corresponds to the bottom-most slice of the 3D panicle.

#### Trait Extraction

In this study, each point cloud was voxelized for volume quantification (Cohen-Or & Kaufman, 1995). The corresponding resulting volume was then used to extract traits of interest, for instance, voxel count and color intensity (Sandhu *et al*. 2019). Also, we calculated the projected surface area. The projected surface area was used to estimate the surface area of the panicle. We first calculated point cloud’s main directions using principal component analysis (PCA) to compute the projected surface area. There were three main directions in a given 3D point cloud. We built 3D coordinate system using the first main direction as Z-axis and the other two directions as the X- and Y-axes. The origin of the system was defined as the lowest point of the point cloud, which was located at the bottom of the panicle. Then, we generated a plane using Y-axis as the norm. After projecting the point cloud of the panicle onto the plane, we calculated the projected surface area as the area of the region enclosed by the boundary of the projected 2D points (Fig. 1b). Afterward, we rotated the plane around the Z-axis and calculated the projected surface area every 5 degrees. In total, we captured 36 projected surface areas. We finally extracted the maximal projected area, the minimal projected area, and the averaged projected area from these results. We used averaged projected area for final analysis and normalization of panicle’s photosynthetic parameters. We also computed the projection area when the plane was perpendicular to the X-axis and Y-axis. Apart from computing the image derived traits (projected surface area, voxel counts, and color intensity) from an entire panicle, we also examined additional traits extracted from local regions of the panicles. We divided the 3D panicle into 10 equal sections along the Z-axis to generate 10 slices. For each slice, we analyzed the corresponding traits (i.e., point count and point color). The analysis of sliced 3D traits enabled us to examine spatial and temporal variation in the development of grains on a particular panicle.

#### Leaf and Panicle photosynthetic measurements

Two LI-6800 (LI-COR) devices were used in parallel to measure leaf and panicle-based gas exchange variables (Fig. 4). All photosynthetic measurements were recorded between 1100-1400 hours. For photosynthetic measurements, the environmental conditions were set as: Relative humidity chamber at 50%, flow rate at 700 µmol s^-1^, chamber pressure at 0.05 kPa, light intensity at 800 µmol m^-2^s^-1^, and reference CO_2_ at 400 µmol mol^-1^. LI-6800 warm-up tests were conducted every time before the actual measurements to control the error rates. To maintain the incident radiation intensity between 800-900 µmol m^-2^ s^-1^ in the greenhouse setting, two adjustable additional LED lights (Vipar Spectra; Model: V300) were used as a source of diffused light. These LED lights included IR (Infrared) LEDs that looked dim/invisible and operated at input voltage 120V and 60Hz frequency. Plants were acclimatized to the artificial light for 15-20 minutes before recording the photosynthetic measurements. One of the LI-6800s was used for measuring gas exchange from the flag leaf, while the other one with an equipped cylinder chamber was used to measure panicle-based photosynthesis on the same plant (Fig. 4). The sensor head of LI-6800 was fitted onto the customized cylindrical chamber having a height and radius of 25.4 and 2.8 cm, respectively, to measure panicle-level incidence light. The bottom end of the cylinder was equipped with a rubber stopper with a small hole facilitating insertion of the panicle stalk. The customized chamber was further sealed with modeling clay each time after inserting the panicle to prevent air leakage. Leaf level measurements were taken after inserting a specified region of a flag leaf into the LI-6800 head; however, panicle-based measurements encompassed the whole panicle. Further, to verify the functioning of our customized chamber, we measured the photosynthetic parameters of young leaves using both traditional chamber and customized chamber (Fig. S6). Unlike panicle, we were not able to completely control the leakage by using customized chamber for taking leaf level photosynthetic measurements. Although no significant differences were observed between values derived from traditional chamber and customized chamber for the leaf measurements, the values from customized chamber were slightly higher due to minimal leakage (Fig. S6). The gas exchange readings from flag leaf and panicle of a particular plant stabilized after 10-15 minutes, after which the setup was used for measuring the next plant. Since LI-6800 measured gas exchange variables were based on per unit area, the surface area of panicles was determined by panicle imaging and then used to normalize the data. The parameters considered for photosynthetic measurements were *A*_*leaf*_ (leaf carbon assimilation), *gsw*_*leaf*_ (leaf stomatal conductance), *E*_*leaf*_ (apparent leaf transpiration rate), *VPD*_*leaf*_ (vapor pressure deficit inside leaf chamber), *A*_*panicle*_ (panicle carbon assimilation), and *E*_*panicle*_ (apparent panicle transpiration rate). The term “apparent” transpiration rate was used in this study to distinguish it from the transpiration rate occurring under natural unenclosed conditions. Furthermore, we calculated water use efficiency (WUE) of leaf (*WUE*_*leaf*_) and panicle (*WUE*_*panicle*_) separately by dividing respective carbon assimilation rate (*A*) with their apparent transpiration rate (*E*).

#### Correlation analysis

We considered data from 3 digital (green pixels proportion, voxel count, and projected panicle area) and four physiological (A_*panicle*_, E_*panicle*_, A_*leaf*_, E_*leaf*_) traits for computing a pairwise Pearson correlation (PCC). Each trait consisted of an observation from three biological replicates under control and HS from accessions TEJ-1 and TEJ-2. PCC between a pair of traits was computed in RStudio v.1.2.5033 platform. We computed PCC separately for TEJ-1 and TEJ-2 at 10 DAF under HS, as the two accessions had a contrasting performance at this time point under HS. The correlation matrix plot and significance level was generated using the ‘chart.Correlation’ function incorporated in the ‘PerformanceAnalytics’ package.

#### Mature seed analysis

To assess effect of moderate HS on mature seeds, we first evaluated only florets marked at the time of fertilization (Dhatt *et al*., 2021). For this, we scored the total number of fully developed and unfilled or completely sterile seeds to calculate percentage fertility. The dehusked mature seeds were used to measure (i) morphometric parameter (length, width), (ii) single grain weight, (iii) percent fertility. Morphometric analysis was performed on 350–1000 marked seeds from 20– 40 plants using *SeedExtractor* (Zhu *et al*., 2021a). Secondly, to have insights into yield-related parameters at a whole plant level, we evaluated all the seeds for percentage fertility and total seed weight per plant.

## Results

### Heat stress induces differential morphological responses in panicles

The purpose of this study was to establish whether multi-view images captured by using PI-Plat could be combined with a novel method for whole panicle gas exchange measurements to follow photosynthetic dynamics during reproductive development. We imposed a moderate HS for 4 or 10 days beginning one day after fertilization (DAF) and measured the photosynthetic response of both foliar and panicle tissue under control (28/23°C; day/night) and HS (36/32°C; day/night). Two rice lines (TEJ-1 and TEJ-2) genetically diverse in their response to HS were compared. From captured images from multiple angles 3D point clouds of panicles were reconstructed to extract the digital traits of a panicle (Fig. 1). The derived digital traits included, projected panicle area (PPA), voxel count (VC), and color intensity (red and green pixels) representing the panicle’s area, volume, and green/red pixel proportion, respectively (Gao et al., 2021; Sandhu et al., 2019). We first used the digital traits to examine whether they could distinguish temporal differences in inflorescence architecture due to HS, and then whether the response differed between TEJ-1 and TEJ-2 (Fig. 2). In TEJ-1, PPA exhibited an increase from 4 to 10 DAF under control conditions, while no significant change was observed under HS (Fig. 2a). An increase in PPA was also observed in TEJ-2 from 4 to 10 DAF under control conditions. However, unlike TEJ-1, TEJ-2 exhibited an increase in PPA from 4 to 10 DAF under HS (Fig. 2a). VC also exhibited a similar trend as PPA in both the genotypes under control and HS (Fig. 2b). For downstream panicle level gas exchange, we decided to use PPA as the normalizing parameter. The PPA and VC for TEJ-2were lower than for TEJ-1 under both temperature conditions, confirming our direct observation of TEJ-2 having a smaller panicle than TEJ-1 (Fig. 2a and 2b).

**Figure 2.**
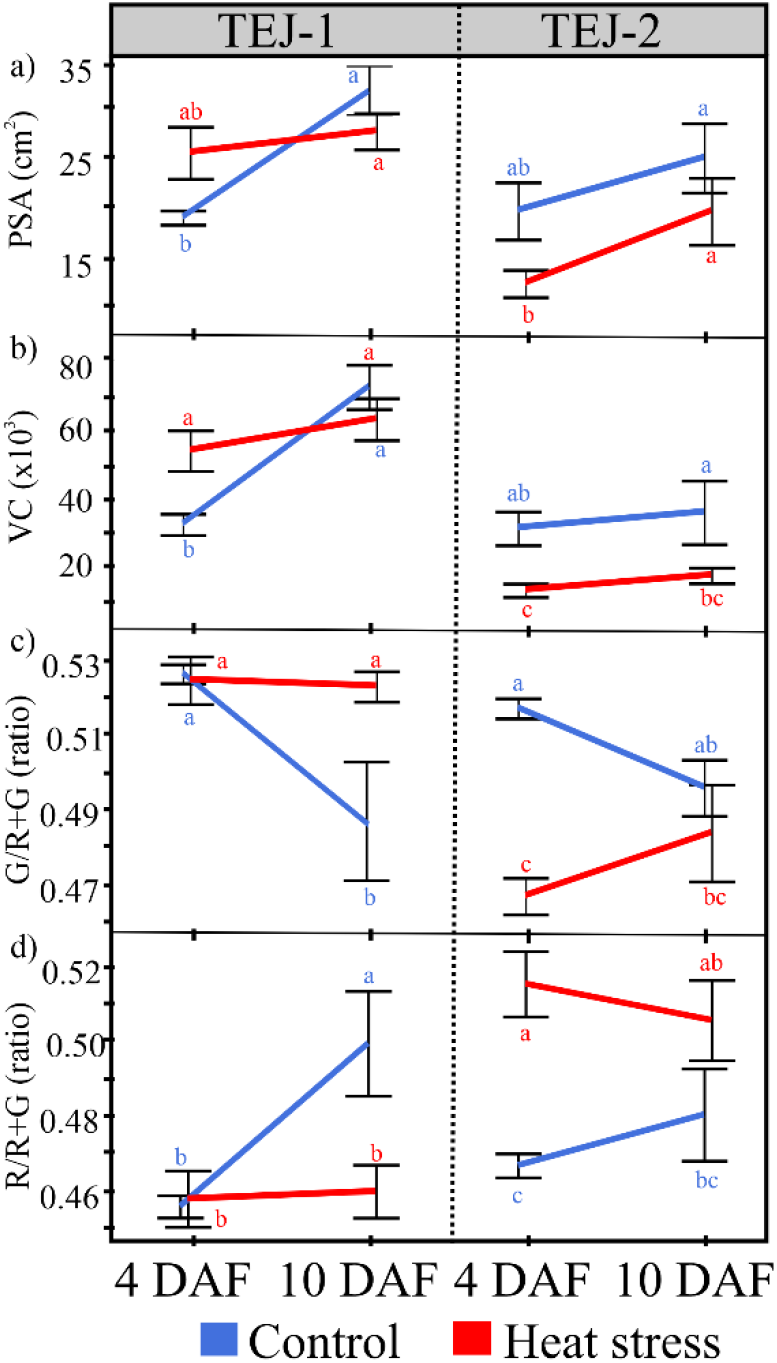
Digital trait analysis from 3D reconstructed panicles of TEJ-1 and TEJ-1 under control (28/23°C; day/night) and HS (36/32°C; day/night). HS was imposed 1 DAF and the traits were measured at 4 DAF and 10 DAF. (a) PPA (Projected panicle area) in cm^2^, (b) VC (Voxel count) representing the point count in a 3D plane, (c) Ratio of green pixels (G) to the sum of red and green pixels (R+G) in a 3D plane, (d) Ratio of red pixels (R) to the sum of red and green pixels (R+G) in a 3D plane. n = 3-4 plants per data point. For statistical analysis, student’s t-test was used to compare respective control and heat stress (α = 0.05). Significant differences are indicated by different letters. Error bars represent ±SE.

### Panicle photosynthetic response to heat stress is dynamic

We next determined whether the gas exchange response of the primary panicle and its corresponding flag leaf varied under the conditions described above (Fig. 3). A standard leaf chamber of the open infra-red gas analyzer was used for the flag leaf. The foliar and non-foliar photosynthetic measurements were conducted the same day as the panicle imaging. For TEJ-1 we observed significantly lower (p < 0.001) stomatal conductance (*gsw*_*leaf*_) for the flag leaf under HS compared to controls at both the time points (4 and 10 DAF) (Fig. S2). In contrast, flag leaf of TEJ-2 exhibited higher *gsw*_*leaf*_ at both 4 and 10 DAF under HS (Fig. S2). Since apparent transpiration rate (*E*) is a function of stomatal conductance, *E*_*leaf*_ also remained significantly lower (p < 0.001) for the TEJ-1 plants grown under HS at both time points compared to controls (Fig. 4a). TEJ-2 plants had higher *E*_*leaf*_ under HS (Fig. 4a). Consistent with stomatal conductance (*gsw*_*leaf*_), recorded carbon assimilation (*A*_*leaf*_) was significantly lower (p <0.001) for TEJ-1 plants under HS at both the time points (Fig. 4a). The carbon assimilation (*A*_*leaf*_) rate of TEJ-2 did not change significantly under HS at 4 and 10 DAF (Fig. 4a). Furthermore, we observed that leaf water use efficiency (*WUE*_*leaf*_) in TEJ-1 was significantly less under HS than control at both 4 and 10 DAF, with a decreasing trend (Fig. S7). In contrast, TEJ-2 exhibited an increasing trend for *WUE*_*leaf*_ in HS and decreasing trend in control from 4 to 10 DAF (Fig. S7). This data suggest that TEJ-1 exhibits greater gas exchange sensitivity in foliar tissue to HS relative to TEJ-2.

**Figure 3.**
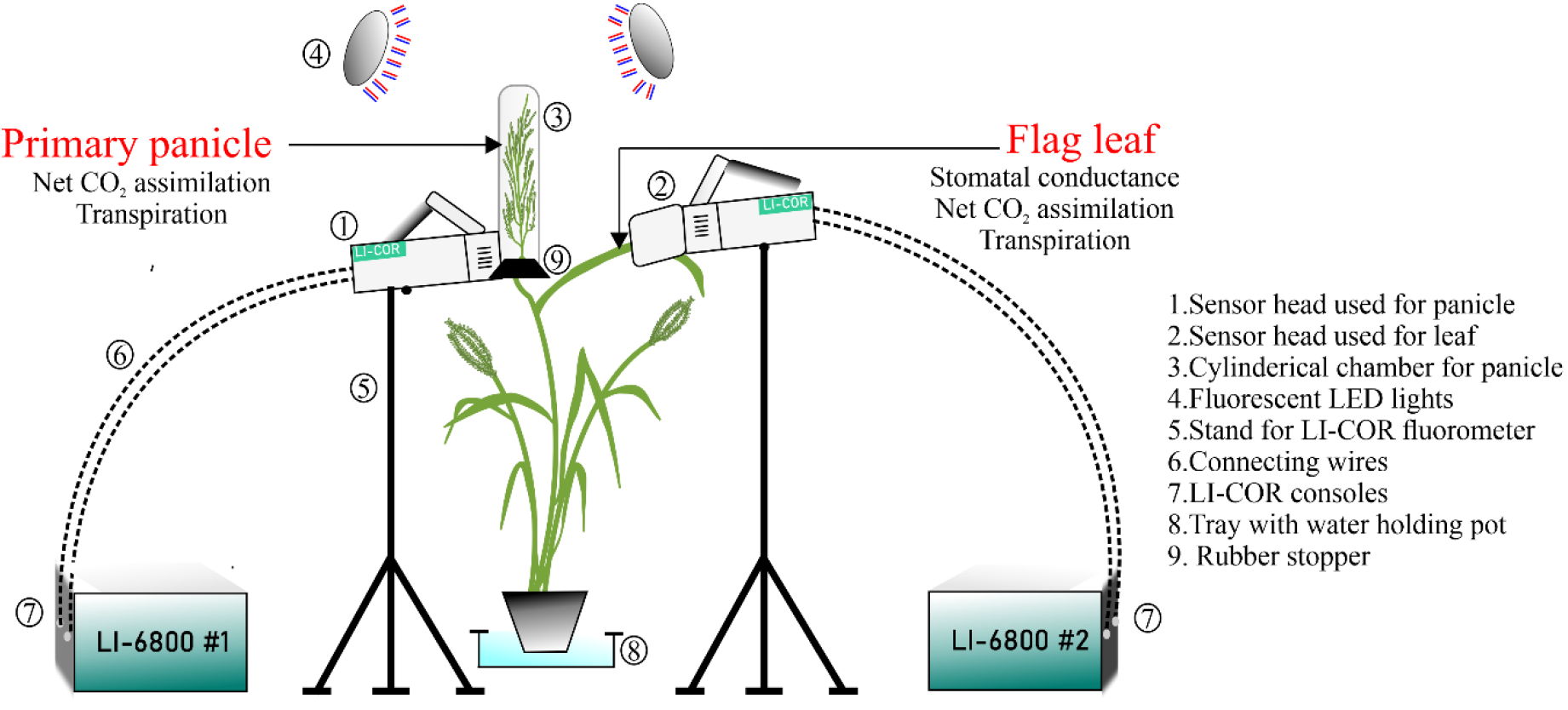
Pictorial representation of the setup used for measuring gas exchange parameters of flag leaf and primary panicle simultaneously using two LI-6800s. The photosynthetic parameters obtained from leaf and panicle that are used for comparative analysis in the study are mentioned in the picture. Numbers represent details of each part of the setup.

**Figure 4.**
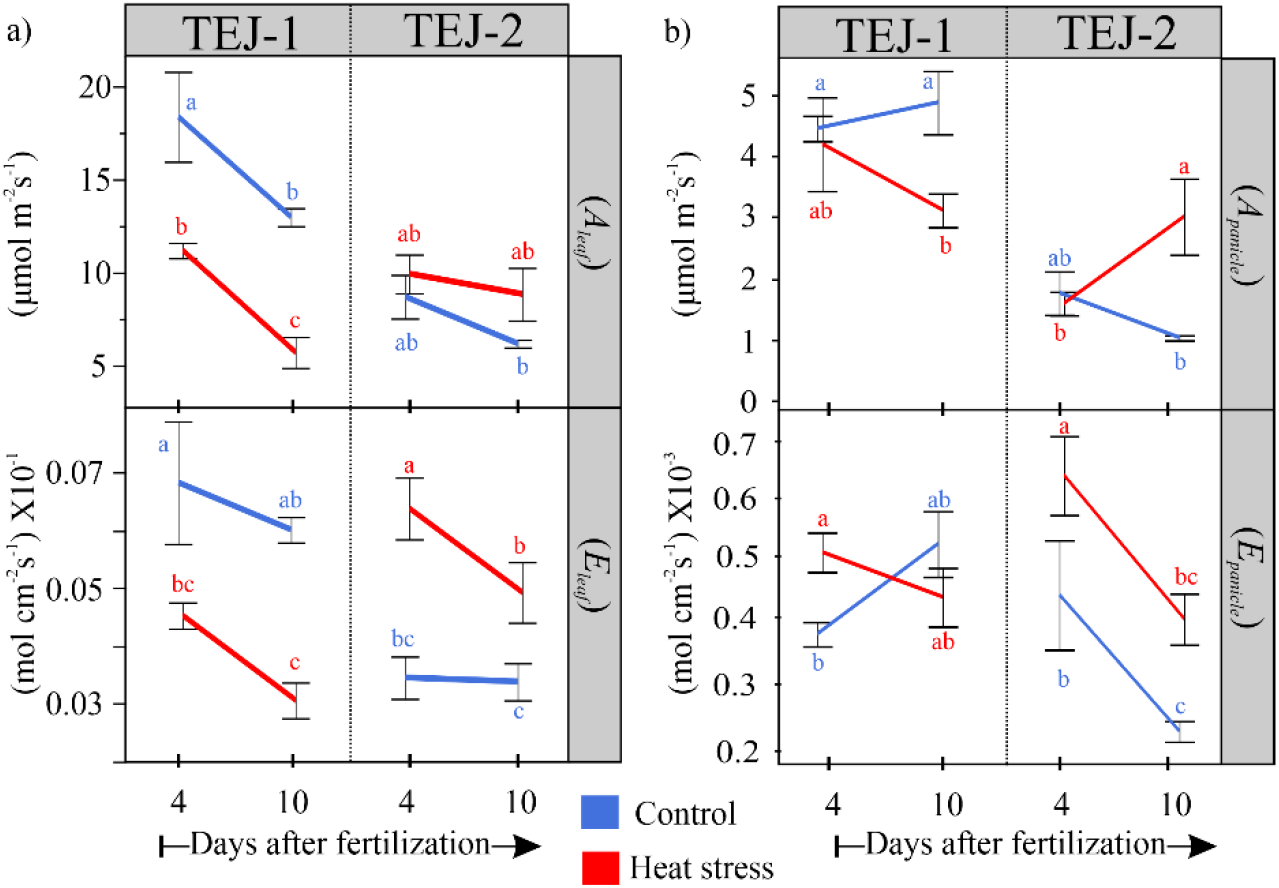
Gas exchange parameters for TEJ-1 and TEJ-2 (a) Flag leaf and (b) Panicle developing under control and heat stress conditions at 4 and 10 DAF (A: Net CO_2_ assimilation; E: Transpiration). N=3-4 plants per data point. For statistical analysis, student’s t-test was used to compare respective control and heat stress values of each of the traits (α = 0.05). Significant differences are indicated by different letters. Error bars represent ±SE. Blue and red color represents control and heat stress, respectively.

We next measured the panicle level photosynthetic response of TEJ-1 and TEJ-2 under HS using a custom-built LI-6800-compatible cylindrical chamber for panicle measurements (Fig. 3). We used PPA for normalizing panicle measurements across genotypes and treatments on a per unit area basis (Fig. 1b). In TEJ-1, there was no difference between *A*_*Panicle*_ under control and HS at 4 DAF (Fig. 4b). However, the *A*_*Panicle*_ was reduced under HS at 10 DAF in TEJ-1. Like TEJ-1, no difference in *A*_*Panicle*_ under control and HS was observed at 4 DAF in TEJ-2 (Fig. 4b). Notably, in TEJ-2, the *A*_*Panicle*_ was higher under HS than control at 10 DAF (Fig. 4b-upper part). The panicle level apparent transpiration rates (*E*_*Panicle*_) were higher under HS than control at 4 DAF in both accessions (Fig. 4b). At 10 DAF, the apparent transpiration rate was similar under HS and control in TEJ-1, and higher under HS than control in TEJ-2 (Fig. 4b-lower part). Additionally, panicle water use efficiency (*WUE*_*panicle*_) of TEJ-1 under HS remained significantly lower than control at both the timepoints (Fig. S8). However, *WUE*_*panicle*_ of TEJ-2 exhibited significant increase at 10 DAF under HS than control (Fig. S7). These photosynthetic measurements indicate that TEJ-1 and TEJ-2 have contrasting responses under HS for *A*_*leaf*_ and *A*_*panicle*_ at 10 DAF. Further, the percent change observed in *A*_*leaf*_ and *A*_*panicle*_ under HS when compared to corresponding controls at 10 DAF (Fig. S3) quantified this genotypic difference. At 10 DAF, *A*_*leaf*_ and *A*_*panicle*_ were reduced by 56% and 26%, respectively, in TEJ-1 under HS compared to their corresponding controls. In contrast, in TEJ-2, *A*_*leaf*_ and *A*_*panicle*_ increased by 57% and 121% respectively, under HS relative to controls (Fig. S3). Collectively, these analyses indicate the potential of our experimental approach involving concurrent measurement of foliar and non-foliar photosynthetic parameters to discern genotypic differences for photosynthetic parameters under heat stress.

Further, we investigated if the panicle-level photosynthetic parameters measured using the cylinder-based chamber can be estimated from the digital traits extracted from the 3D reconstructed panicles. For this we extracted the pixel color intensities from 3D-recontructed panicles to differentiate their response to HS. The 4 and 10 DAF measurements correspond to the active grain filling phase when the panicle is predominantly green. Since green (G) pixel intensity can be used as a proxy for panicle surface chlorophyll content, we estimated the proportion of green pixels to the sum of red and green pixels [G/(R+G)] to determine changes in response to HS. Under control conditions, TEJ-1 exhibited a decline in green pixel proportion from 4 to 10 DAF (Fig. 2c). While under HS, no significant decline was observed from 4 to 10 DAF in green pixel ratio in TEJ-1 (Fig. 2c). The proportion of green pixels decreased from 4 to 10 DAF in TEJ-2 under control conditions (Fig. 2c). These observations did not explain the change or lack of change in photosynthetic parameters for both genotypes under control conditions. However, the proportion of green pixels increased from 4 to 10 DAF in TEJ-2 under HS (Fig.2c). This observation was consistent with the striking increase observed in *A*_*panicle*_ in TEJ-2 at 10 DAF under HS (Fig. 4b-upper part). As panicle approaches maturity, pixels are expected to shift towards R. Therefore, we also analyzed the proportion of red pixels to the sum of red and green pixels [R/(R+G)]. In TEJ-1, the proportion of red pixels increased from 4 to 10 DAF under control conditions, while it remained similar between 4 and 10 DAF under HS (Fig. 2d). TEJ-2 also exhibited a similar trend as TEJ-1 for red pixels proportion under control conditions (Fig. 2d). However, the red pixel proportion was higher in TEJ-2 than TEJ-1 under HS at both time points (Fig. 2d). Based on our analysis, whole panicle level G pixel proportion does not correspond well with panicle gas exchange measurements.

### Digital slicing of reconstructed panicles captures panicle level spatial variation

The observed inconsistency between photosynthetic parameters and green pixel proportion, promoted us to further examine the pixel color intensities by accounting for spatial variability along the panicle length due to variable developmental stage of the seeds, resulting from asynchronous fertilization. Therefore, we divided the 3D reconstructed panicle into ten equal slices. Digital traits were obtained for individual slices (Fig. 1) and compared between control and HS for each genotype (Fig. 5). We performed spatial analysis for VC and green pixel proportion [G/(R+G)] for both genotypes (Fig. 5 and S5). In TEJ-1, a gradient in green pixel proportion was observed from top slices (slices 1-4) having higher green pixel proportion than lower slices (slices 5-10) at 4 DAF under control conditions (Fig. 5a). By the 10 DAF time point, the top slices (slices 1-4) had reduced green pixel proportion and the bottom slices (slices 5-10) had increased green pixel proportion under control conditions in TEJ-1 (Fig. 5a). Unlike control conditions, a gradient in green pixel proportion was observed with middle slices (slice 4-7) having higher proportion, followed by bottom slices (slices 8-10), and then the top slices (slices 1-3) at 4 DAF under HS in TEJ-1 (Fig. 5a). The green pixel proportion of upper slices (slices 1-4) increased at 10 DAF compared with 4 DAF, whereas they were lower for most of the bottom slices (slices 5-10; except slice 7) under HS in TEJ-1 (Fig. 5a). TEJ-2 also had a gradient in green pixel proportion under control conditions at 4 DAF with the top slices (slices 1-4) having higher green pixel proportion than the bottom slices (slices 5-10) (Fig. 5b). At 10 DAF, the top slices (slices 1-4) had a reduced green pixel proportion, while the bottom slices (slices 5-10) had similar green pixel proportions as those of 4 DAF under control conditions in TEJ-2 (Fig. 5b). A notable feature of the TEJ-2 under HS was its ability to largely maintain a higher green pixel proportion for the bottom slices (slices 7-10) at 4 and 10 DAF relative to control values (Fig. 5b). At 10 DAF in TEJ-2, the green pixel proportion for top slices (slices 1-6) increased slightly compared to 4 DAF under HS (Fig. 5b). Collectively, variations in the green pixel proportion pattern obtained from slicing of 3D panicles illustrates the spatial heterogeneity among the florets and its transition with progression of both development and stress duration.

**Figure 5.**
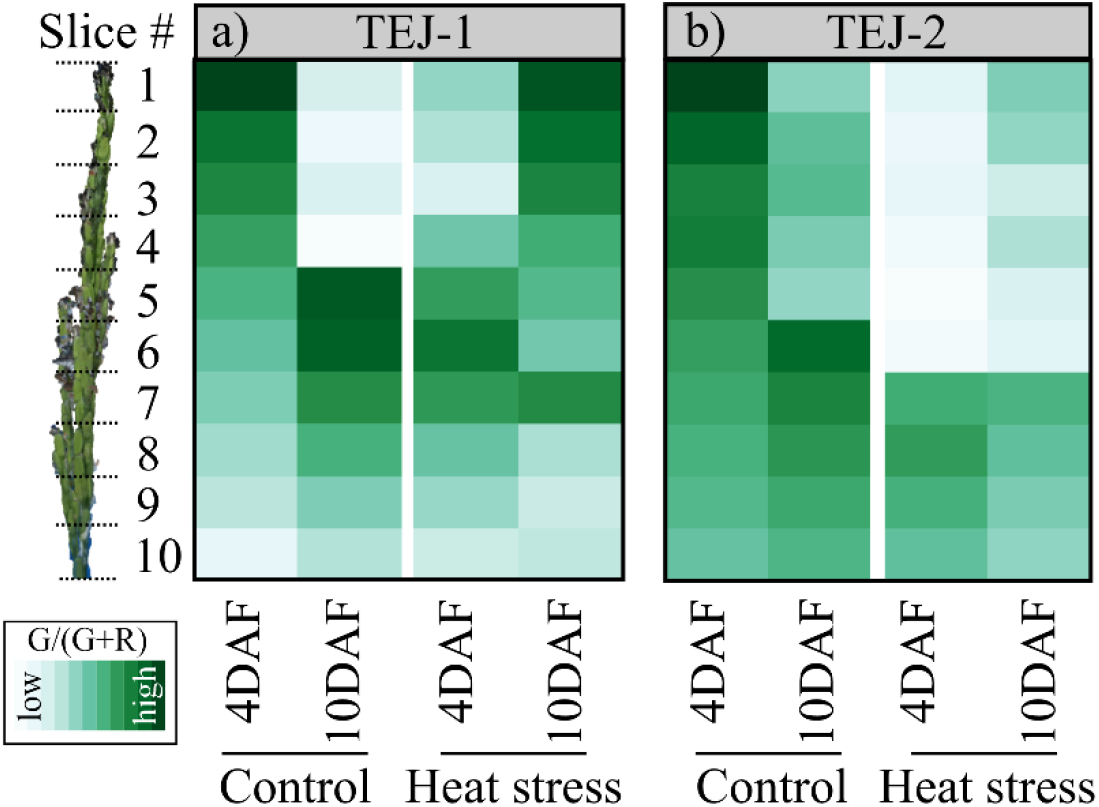
Shifts in green pixel intensity resolved into 3D slices using the panicle point cloud (a) TEJ-1 and (b) TEJ-2. Progression of color from white to green in the heat map represents increase in green pixel intensity, which is a proxy for chlorophyll content of panicle surface. N=3-4 plants per data point. Respective values from each slice of all the replicates were averaged to make the final heat map. Control and heat stress values of green pixel intensity for the genotypes are on same scale to show the temporal and spatial changes.

### Correlations between digital traits and photosynthetic measurements vary with genotypes

We next examined the relationship among 3D reconstruction-derived features and photosynthetic parameters for the genotypic responses to HS at 10 DAF. We selected the 10 DAF for this analysis as we observed the most significant genotypic contrast at this time point under HS. We used the digital traits (PPA, VC, and G) and photosynthetic measurements (A_*panicle*_, E_*panicle*_, A_*leaf*_, E_*leaf*_) to perform pairwise correlation analysis separately for TEJ-1 and TEJ-2 under HS. The derived digital traits, i.e., PPA, VC, and green pixel proportion, showed a strong positive correlation among themselves and a negative correlation with A_*leaf*_, A_*panicle*_, and E_*leaf*_ in both genotypes (Fig. 6, green boxes). Further, the correlation between some of the examined parameters exhibited contrasting values in TEJ-1 and TEJ-2 (Fig. 6, blue boxes). Although these correlation values between particular digital traits and photosynthetic parameters were not statistically significant, they still suggest a divergent reponse for TEJ-1 and TEJ-2 under HS. For instance, in TEJ-1, the correlation of *A*_*panicle*_ with PPA, VC, and G was -0.42, -0.80, and -0.66, respectively (Fig. 6a, blue boxes), while in TEJ-2, the correlation of *A*_*panicle*_ with PPA, VC, and G were +0.43, +0.70, and +0.69, respectively (Fig. 6b, blue boxes). These results suggest that despite having a larger panicle size and higher pixel count under HS, TEJ-1 does not exhibit an increase in *A*_*panicle*_, resulting in negative correlation values. In TEJ-2, *A*_*panicle*_ increases along with PPA, VC, and G under HS, resulting in a positive correlation. Further, the correlation between *A*_*panicle*_ and *A*_*leaf*_ in TEJ-1 and TEJ-2 was +0.88 and -0.68, respectively (Fig. 6, blue box). These results suggest that in TEJ-1, both *A*_*panicle*_ and *A*_*leaf*_ are decreasing under HS, leading to a positive correlation value (Fig. 2 and 6), while TEJ-2 has higher *A*_*panicle*_ and more stable *A*_*leaf*_ under HS, resulting in negative correlation (Fig. 2 and 6).

**Figure 6.**
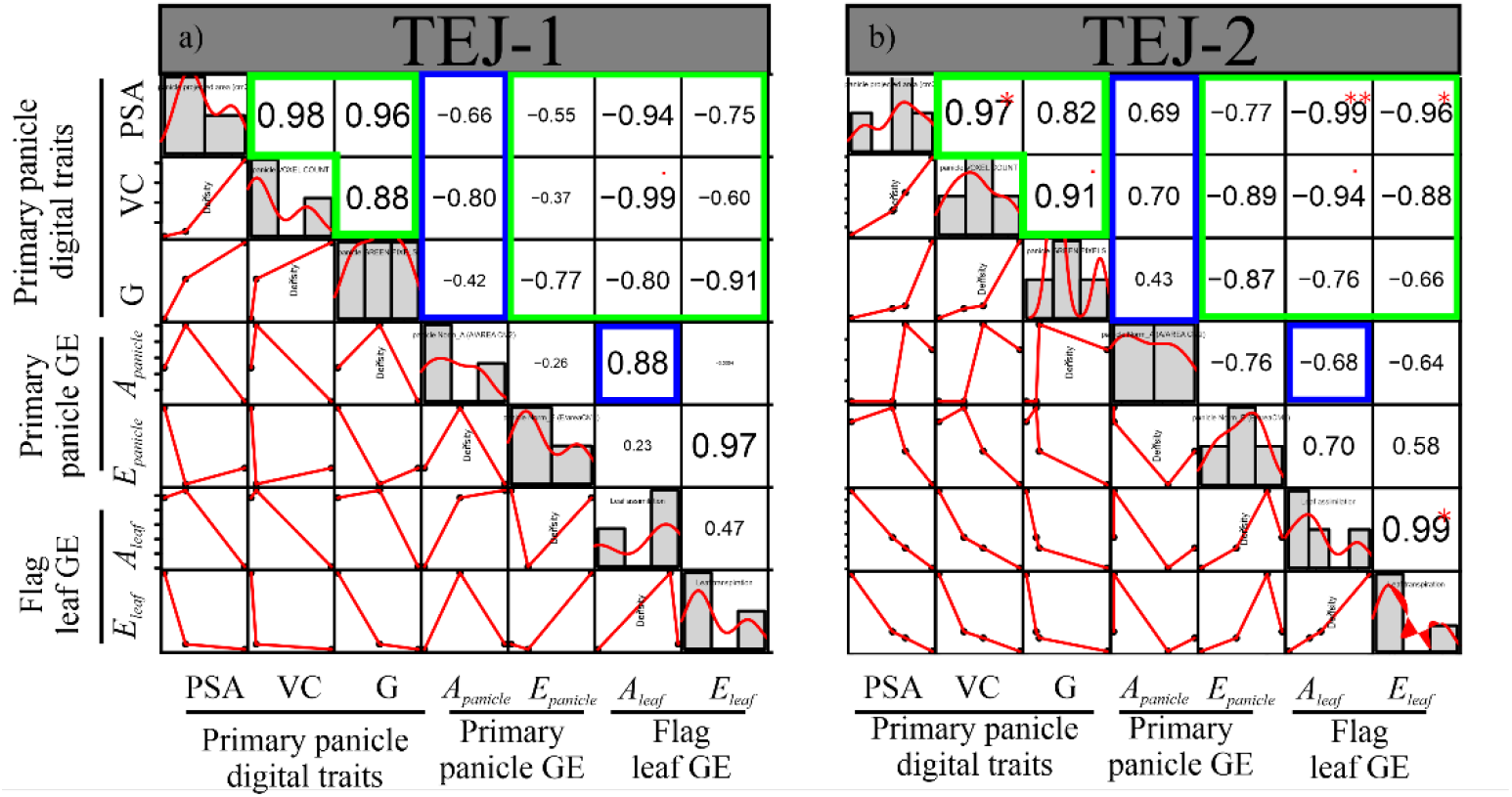
Correlation of primary panicle digital traits, primary panicle gas exchange (GE) parameters and flag leaf GE parameters at 10 DAF under HS in (a) TEJ-1 and (b) TEJ-2. Histograms and red lines represent each trait’s distribution. Green colored boxes indicate similar type of correlation in TEJ-1 and TEJ-2 for the respective traits. Blue colored boxes represent contrasting correlation values in TEJ-1 and TEJ-2 for underlying traits. PSA: projected surface area, VC: voxel count, G: proportion of green pixel intensity, *A*_*panicle*_: Net CO_2_ assimilation of primary panicle, *E*_*panicle*_: Transpiration of primary panicle, *A*_*leaf*_: Net CO_2_ assimilation of flag leaf, *E*_*leaf*_: Transpiration of flag leaf, GE: gas exchange. (**, p < 0.01; * p < 0.05.)

### Analysis of mature grain parameters of TEJ-1 and TEJ-2 under HS

The digital traits from 3D reconstructed panicle and photosynthetic measurements indicate that TEJ-1 and TEJ-2 have differential response to HS. We next asked if these observed differences at early seed development stages translate to differences in grain traits at maturity. For this, we imposed short (HS-I; 2-4 DAF) and long (HS-II; 2-10 DAF) duration HS and measured seed length, width, weight, and fertility (Fig. S4a). Mature grain parameters of TEJ-1 and TEJ-2 did not differ significantly different between control and HS-I, except for fertility (%), which was higher in TEJ-2 after heat treatment (Fig. S4b). Under HS-II, fertility was significantly reduced in TEJ-1 but not in TEJ-2 (Fig. S4b). Seed length was not affected in TEJ-1 but increased under HS-II in TEJ-2. A significant reduction in seed weight and width of marked seeds on the primary panicles was observed for both TEJ-1 and 2 at HS-II compared to respective controls (Fig. S4b). The results indicate that TEJ-1 and 2 exhibit differential tolerance to the longer duration heat stress (HS-II) for marked seeds. At the whole plant level, the fertility and per plant grain weight were reduced due to HS-I and HS-II in TEJ-1 compared to its control (Fig. 7a). However, these two parameters were not affected for both heat treatments in TEJ-2. The whole plant level seed trait data suggests that TEJ-2 exhibits greater heat tolerance even for seeds that were fertilized under heat stress compared to TEJ-1. The marked seeds are distinct from whole plant level seeds as they are derived from fertilization events that occur before the imposition of HS treatments.

**Figure 7.**
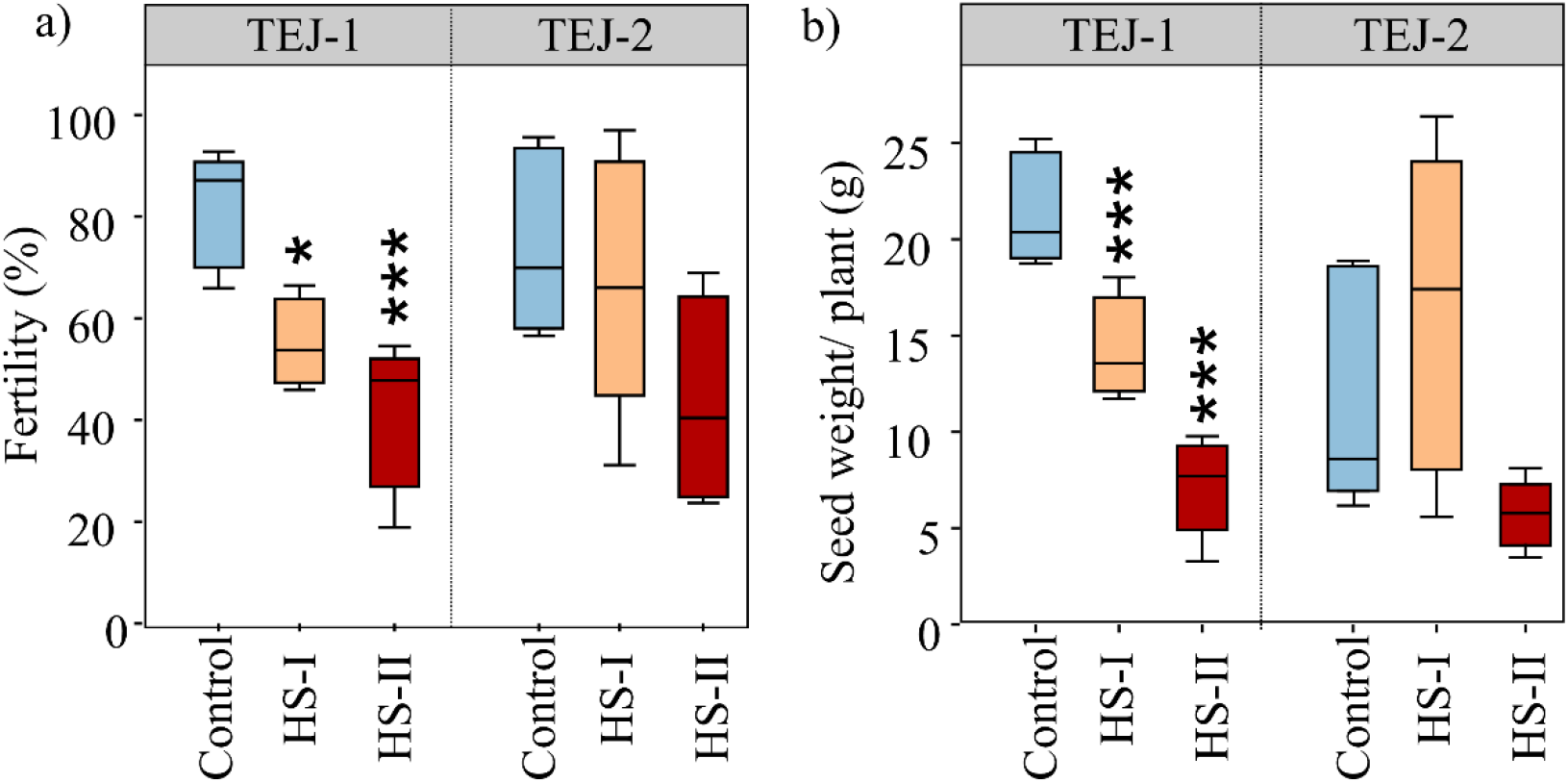
Impact of heat stress on mature seeds at whole plant level in TEJ-1 and TEJ-2 developing under control and heat stress (HS) conditions during grain filling. HS-I and HS-II refer to the duration of imposed HS i.e., 1-4 DAF (HS-I) and 1-10 DAF (HS-II). a) Quantification of spikelet fertility (%) and b) seed weight in grams at the whole plant level evaluated at time of physiological maturity. Box plots show the median and the upper quartiles and black dots signify outliers (5th/95th percentile). N= 1500-3500 seeds from 4-6 plants per data point. For statistics, t-test was used to compare heat stressed mature seeds with respective controls (***, p < 0.001; **, p < 0.01; * p < 0.05.)

## Discussion

It is likely that the negative effects of HS on seed development results partially from disturbance in photosynthesis not only in foliar tissues, but also in non-foliar tissues, as well from the dynamic interactions between these two photosynthate sources. To explore these questions, we developed and tested a novel and more precise method to non-destructively measure rice panicle photosynthetic parameters. Our hypothesis was that this approach, combined with concurrent foliar measurements by traditional methods, would enhance our understanding of the photosynthetic response to HS. Further, we postulated that this method could uncover differences between rice lines that differ in their HS response during reproductive development. Such comparative analyses could eventually help explain why grain fill in some rice accessions is less affected by HS than others. We determined the relative rates of gas exchange between flag leaf and panicle under HS during grain filling stage, the effect of altered carbon fixation (of flag leaf and panicle) due to HS on the final grain yield parameters and distinguish the differential physiological response of two genotypes under HS. In addition to photosynthetic measurements, we also assessed panicle level digital traits to track developmental dynamics along the panicle length under control and HS conditions. For this, we digitally partitioned the 3D reconstructed panicle into ten equal slices and extracted digital traits for each slice. The spatial perspective of the 3D reconstructed panicle enabled us to discern differences between TEJ-1 and TEJ-2 heat stress response at greater resolution (Fig. 5). The analysis of voxel count (VC) and projected panicle area (PPA) from the whole panicles indicated an increasing trend from 4 to 10 DAF in both genotypes under optimal conditions (Fig. 2a & 2b). The spatiotemporal characterization of the panicle slices was able to differentiate responses of the two genotypes that were not evident from whole panicle traits. For instance, whole 3D panicle of TEJ-1 under HS did not exhibit a significant change in the green pixel proportion from 4 to 10 DAF (Fig. 2c). However, sliced 3D panicle results indicate that green pixel proportion of lower (proximal) slices was higher at 4 DAF whereas upper (distal) slices were higher at 10 DAF (Fig. 5a). For TEJ-2, we observed more stable green pixel spatial profile when comparing the 4 and 10 DAF under HS. TEJ-2 slicing results show that the proximal panicle slices (slices 7-10) do not exhibit a drop in the green pixel intensity at 10 DAF under HS (Fig. 5b). This is in contrast with the proximal slices (8-10) in the TEJ-1 at 10 DAF. It is plausible that observed increase in *A*_*panicle*_ at 10 DAF under HS in TEJ-2 could be primarily due to proximal spikelets that “stay green” for longer duration. Alternatively, the panicle architecture of TEJ-2 may be different from TEJ-1 in maintaining growth in proximal part, reflected in largely stable values across time and treatments.

The digital traits derived from 3D reconstructed panicles were able to detect variations in developmental progression of the two genotypes under HS. Since developing grain acts as the active sink tissue, the progression in grain development depends upon accumulation and utilization of the photoassimilates. To examine the source-sink relationship and its effect on grain development, we measured photosynthetic parameters for the flag leaf and primary panicle simultaneously. Apart from the major photosynthetic parameters impacting carbon fixation, parameters like vapor pressure deficit (*VPD*) are known to increase under HS, and hence are a factor for consideration (De Boeck *et al*., 2010; Grossiord *et al*., 2020). Our results show a higher leaf *VPD* for the plants exposed to HS, indicating a greater leaf-atmosphere diffusion gradient (Fig.S2). At higher *VPD*, plants tend to lose more water and trigger stomatal closure to maintain plant water status under limited water conditions (De Boeck *et al*., 2010; Grossiord *et al*., 2020; Moore *et al*., 2021). However, if water availability and *VPD* are not restrictive factors, high temperature can induce guard cell expansion which facilitates stomatal opening to trigger evaporative cooling of the leaf (Faralli *et al*. 2019; Tricker *et al*. 2018; Kostaki *et al*., 2020). The two genotypes in this study showed a contrasting response in foliar gas exchange parameters on exposure to HS under similar growth conditions, including water availability and *VPD* (Fig. 4 and S2). For instance, reduction in leaf stomatal conductance, apparent transpiration rate, and carbon assimilation was observed in TEJ-1 under HS even though plants were growing in well-watered conditions. In contrast, TEJ-2 maintains higher apparent transpiration rate, stomatal conductance, and carbon assimilation under longer duration HS, suggesting that there may be a temperature dependent or independent stomatal response difference between the two genotypes. This could be due to genotypic variation in biomechanical elasticity of the guard cell complex. Alternatively, TEJ-1 may lack the hydraulic structure to sustain water movement under high *VPD* conditions, resulting in differential ABA accumulation in the guard cells.

The non-foliar, panicle-based photosynthetic measurements indicated that net CO_2_ assimilation (*A*_*panicle*_) for both genotypes was similar between optimal and HS conditions at 4 DAF (Fig. 4b). However, *A*_*panicle*_ exhibited a contrasting response in TEJ-1 and TEJ-2 under HS at 10 DAF. TEJ-1 showed a decline and TEJ-2 showed an increase in *A*_*panicle*_ under HS compared to their respective controls at 10 DAF (Fig. 4b). Notably, the apparent transpiration rate for the TEJ-2 declined under HS but the *A*_*panicle*_ increased for 10 DAF panicles. Therefore, estimated WUE for TEJ-2 was also significantly higher than the optimal conditions at 10 DAF (Fig. S8). This decoupling of *A*_*panicle*_ from the apparent transpiration rate in TEJ-2 under HS is intriguing as it likely promotes carbon assimilation rather than evaporative cooling of the panicle.

The photosynthetic parameters measured for two genotypes were consistent with plant-level grain parameters. For instance, TEJ-1, which exhibited a decline in assimilation rate (*A*_*panicle*_ and *A*_*leaf*_) measured during the grain filling stage, also had significantly reduced mature grain weight and fertility parameters (Fig. 4 and 7). TEJ-2 had an enhanced assimilation rate (*A*_*panicle*_ and *A*_*leaf*_) under HS at 10 DAF and showed no significant change in mature grain weight and fertility parameters at whole-plant level (Fig. 4 and 7). In TEJ-1, there was a greater percent decrease in *A*_*leaf*_ (−57%) than in *A*_*panicle*_ (−26%) at 10 DAF under HS as compared to respective controls. In contrast, in TEJ-2 the percent increase in *A*_*leaf*_ (57%) was considerably less than in *A*_*panicle*_ (121%) in response to HS relative to control values. The higher *A*_*panicle*_ for TEJ-2 under HS at 10 DAF is also consistent with the more stable spatial profile of TEJ-2 for green pixel proportion under HS relative to TEJ-1, especially in the proximal end of panicles.

### Conclusion

This work shows the potential value of combining foliar and non-foliar physiological measurements to examine dynamic heat stress response in rice, and to identify genotypic differences in this response. By measuring temporal dynamics along the panicle length, we were also able to discern spatial differences under heat stress. This improved non-destructive approach combines 3D imaging, photosynthetic measurements, and grain physiology, and could be used to gain a spatiotemporal perspective on multiple stress responses and in a variety of cereal species bearing compact inflorescences.

## Acknowledgements

We would like to thank Prof. Timothy Arkebauer at University of Nebraska-Lincoln and Dr. Harel Bacher at Cornell University for their suggestions to improve the manuscript. We would also like to thank Jeremy Hiller for helping in trouble-shooting the technical issues while performing experiments with the LiCor instrument.

## Author contributions

HW, BKD, and JSD wrote the manuscript with contributions from all authors. JSD, BKD, and PP performed the experiments and analyzed the results. JH designed and built the customized cylinder. JSD and BKD calibrated and tested the customized cylinder. TG and HY performed the image analysis. TA and PAS critically reviewed and analyzed the work. All authors read and approved the manuscript.

## Supporting information

**Figure S1.**
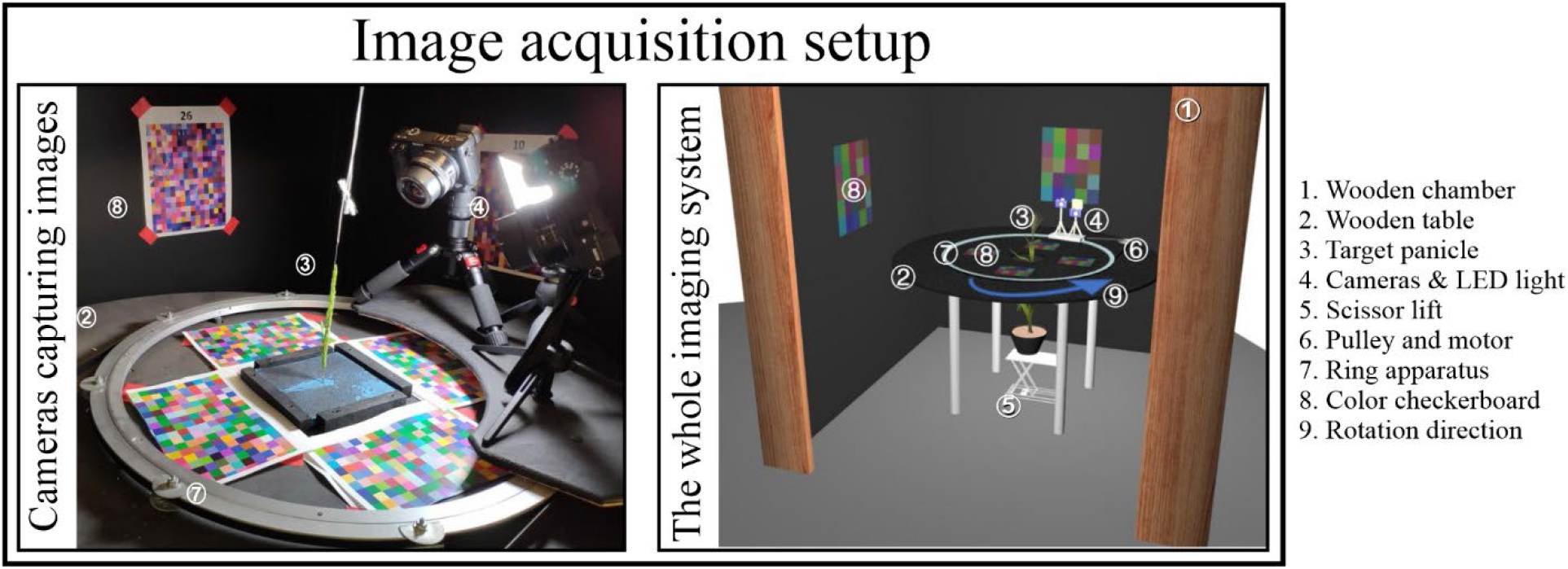
Image acquisition setup using PI-Plat imaging platform.

**Figure S2.**
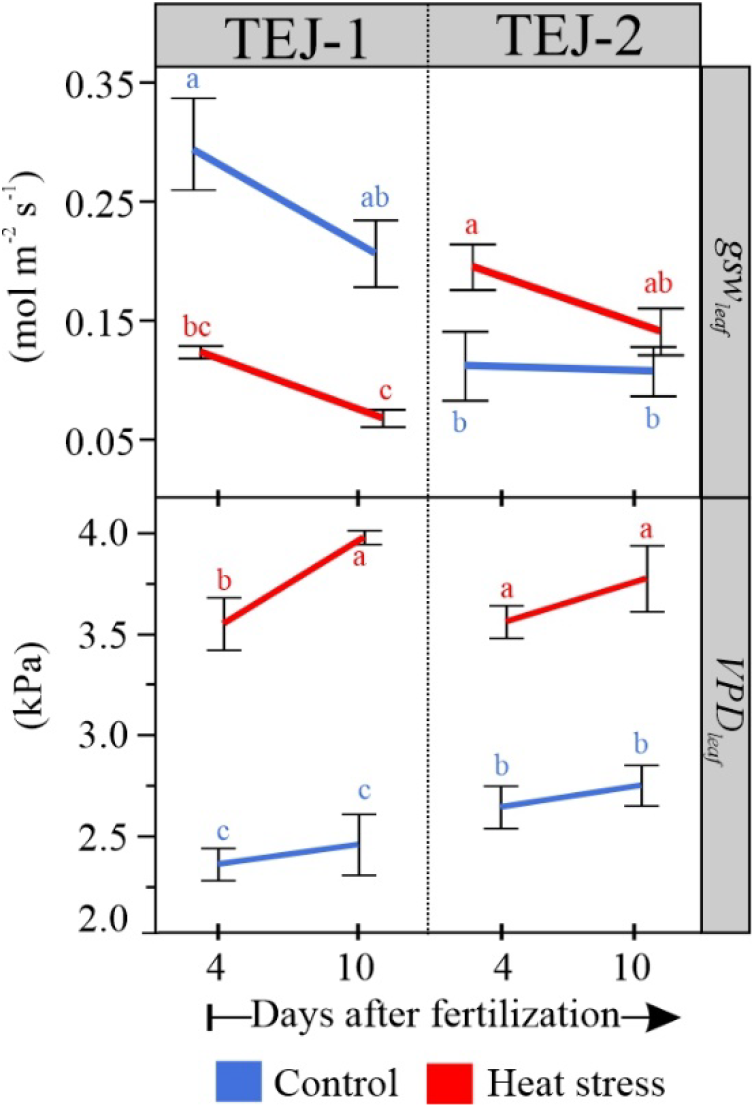
Stomatal conductance (*gsw*) and vapor pressure deficit (*VPD*) of flag leaf of TEJ-1 and TEJ-2 developing under control and heat stress conditions at 4 and 10 DAF.

**Figure S3.**
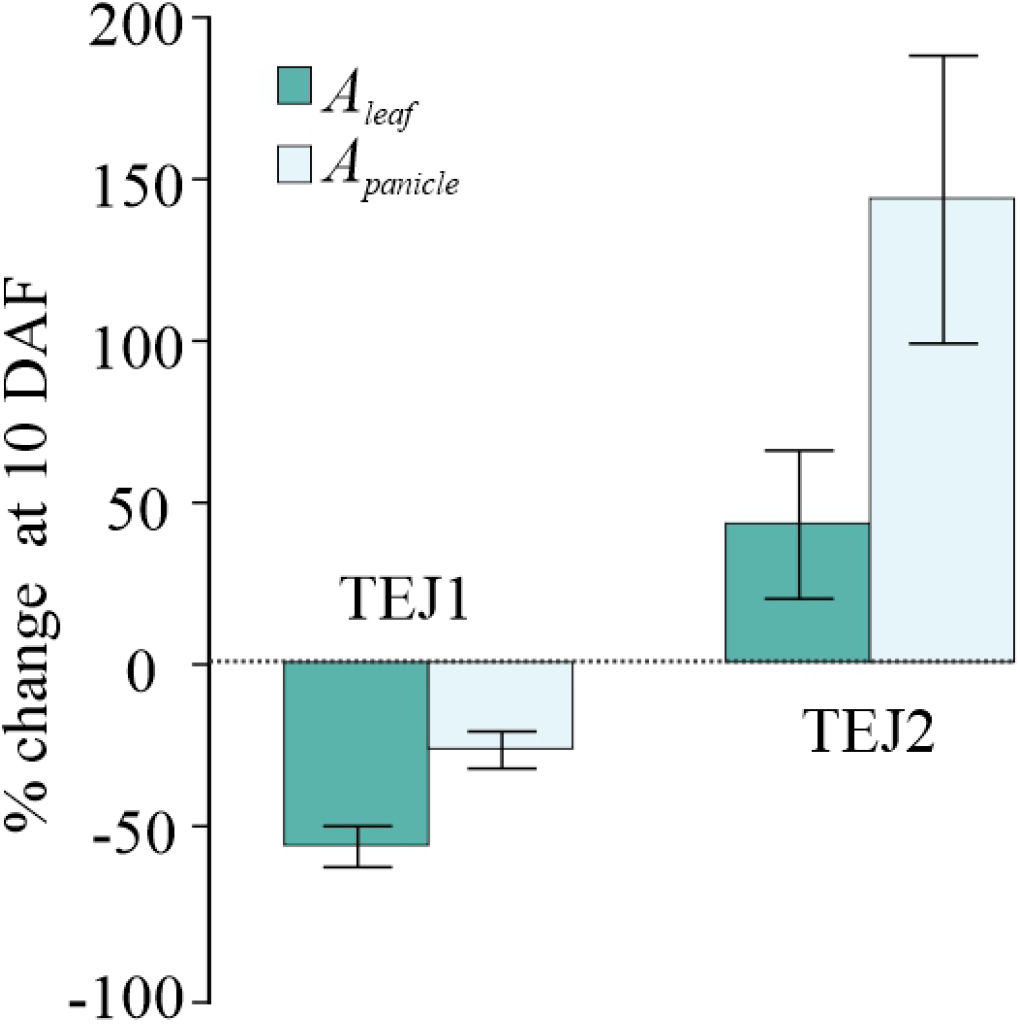
Percent change in A_*leaf*_ and A_*panicle*_ at 10 DAF under HS as compared to respective control values in TEJ-1 and TEJ-2. Error bars represent ±SE.

**Figure S4.**
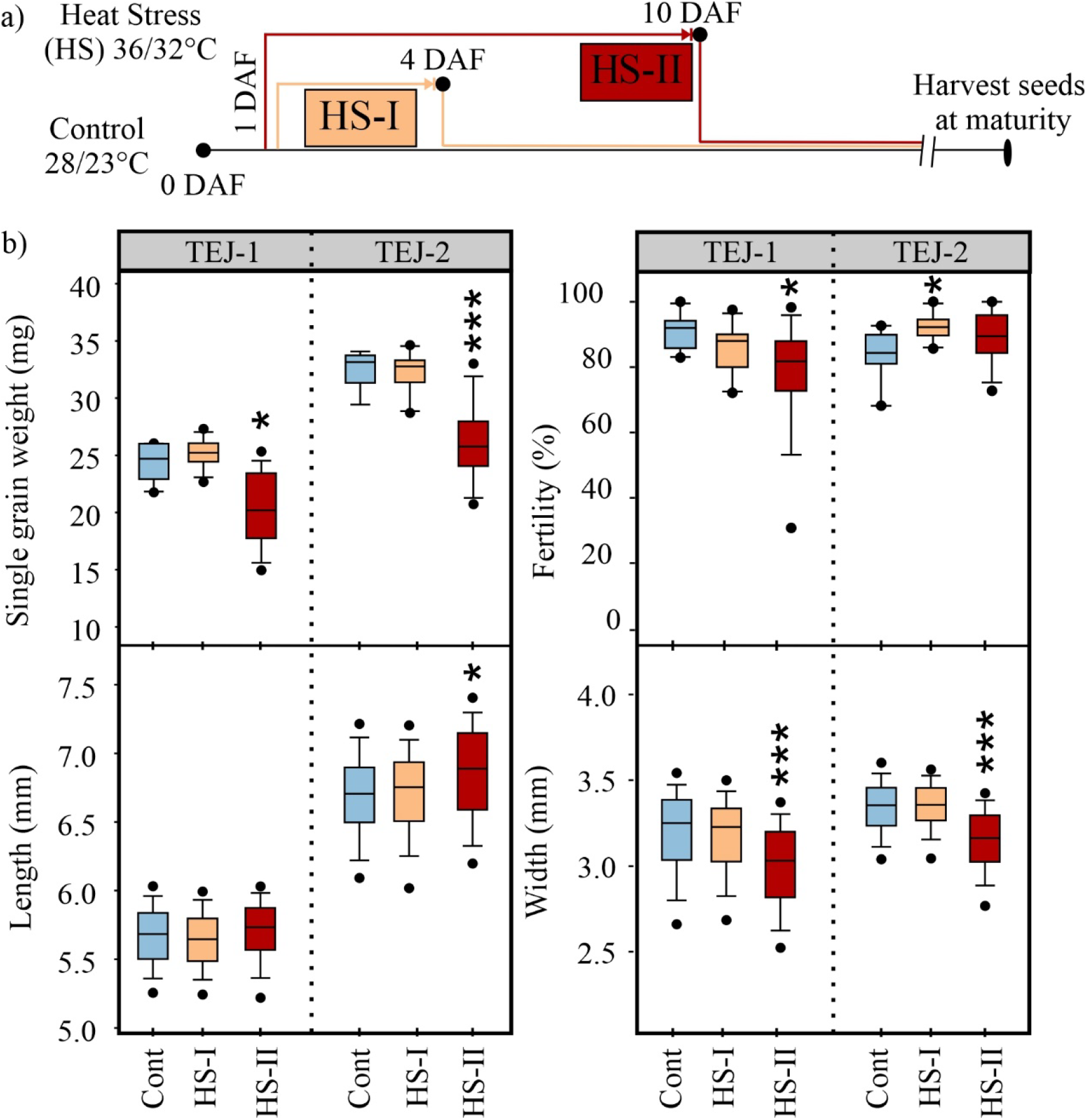
Quantification of single grain weight (mg), spikelet fertility (%), grain length (mm), and grain width (mm) from marked seeds evaluated at the time of physiological maturity.

**Figure S5.**
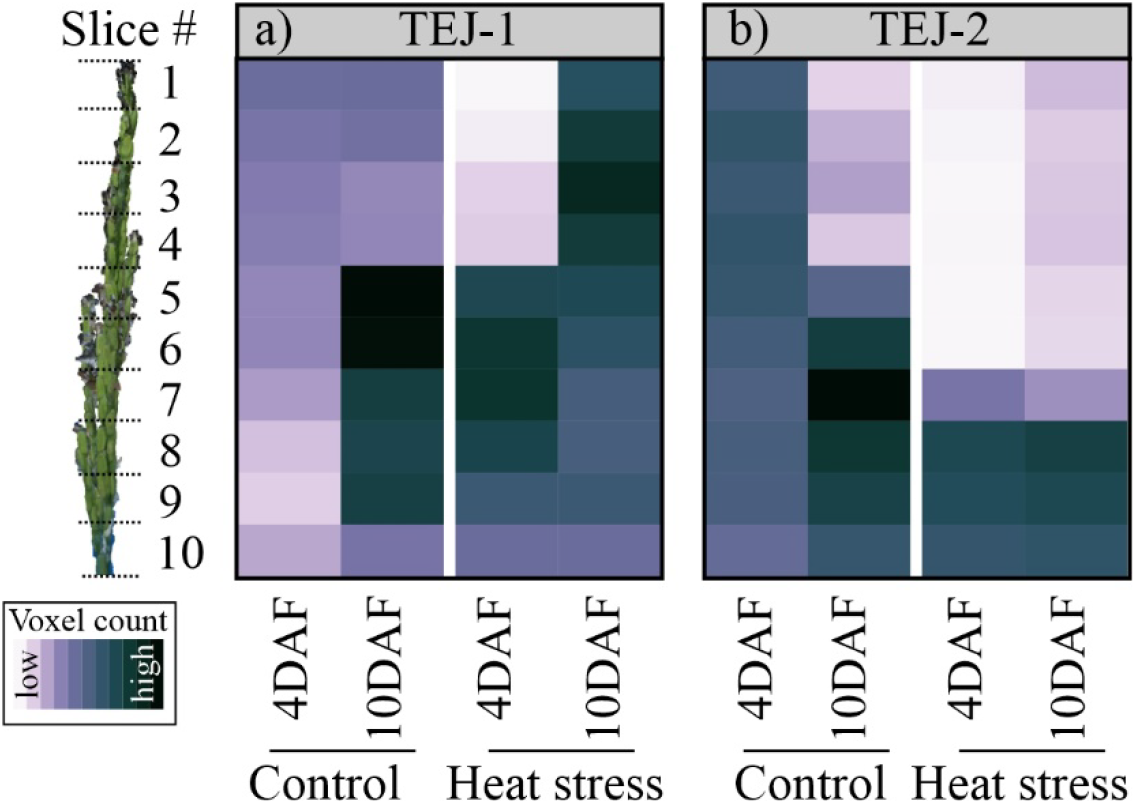
Shift in voxel count resolved into 3D slices using the panicle point cloud (a) TEJ-1 and (b) TEJ-2.

**Figure S6.**
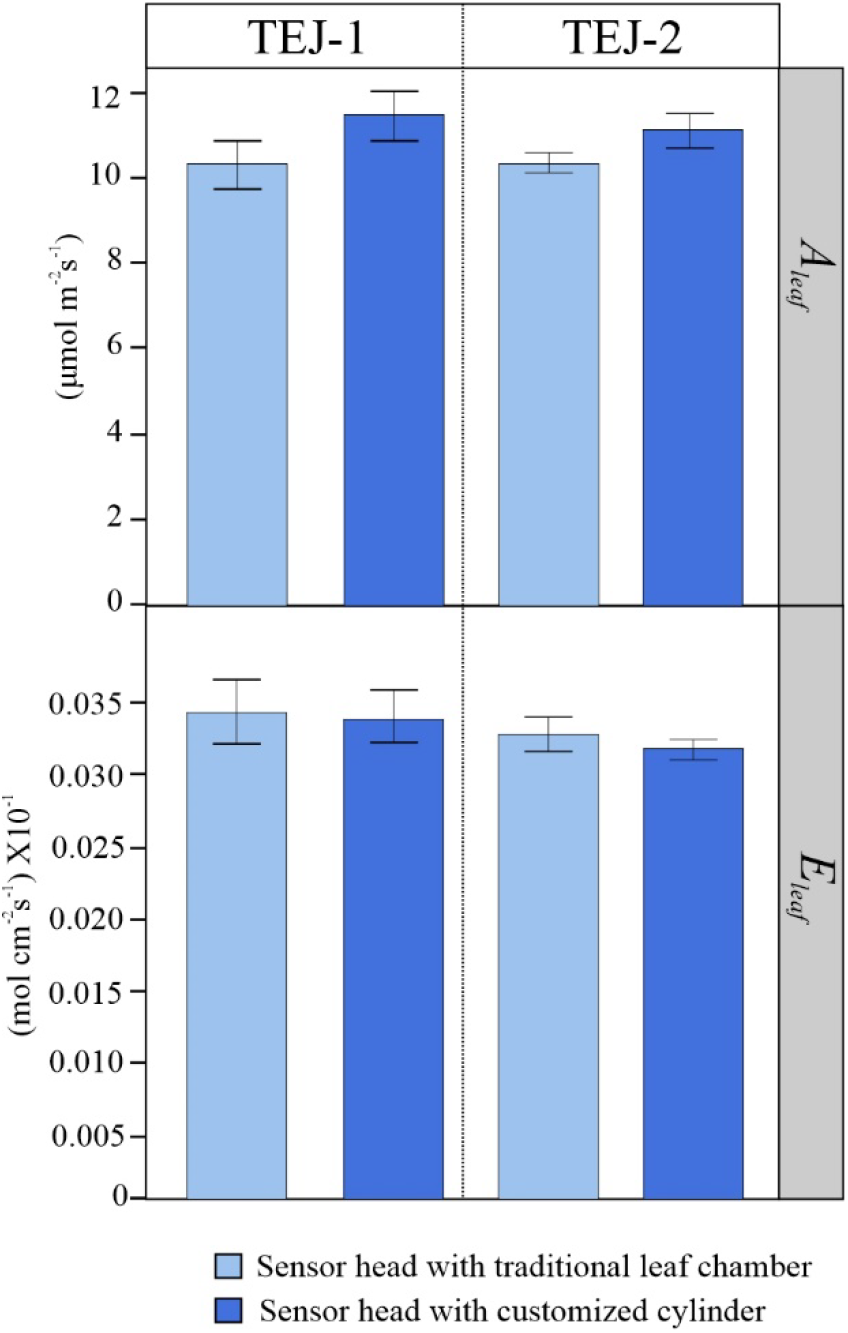
Measurements of Aleaf and Eleaf from randomly selected young green leaf (not flag leaf) of TEJ-1 and TEJ-2 plants using sensor head equipped with traditional leaf chamber (light blue) and customized cylinder (dark blue) under control temperature conditions.

**Figure S7.**
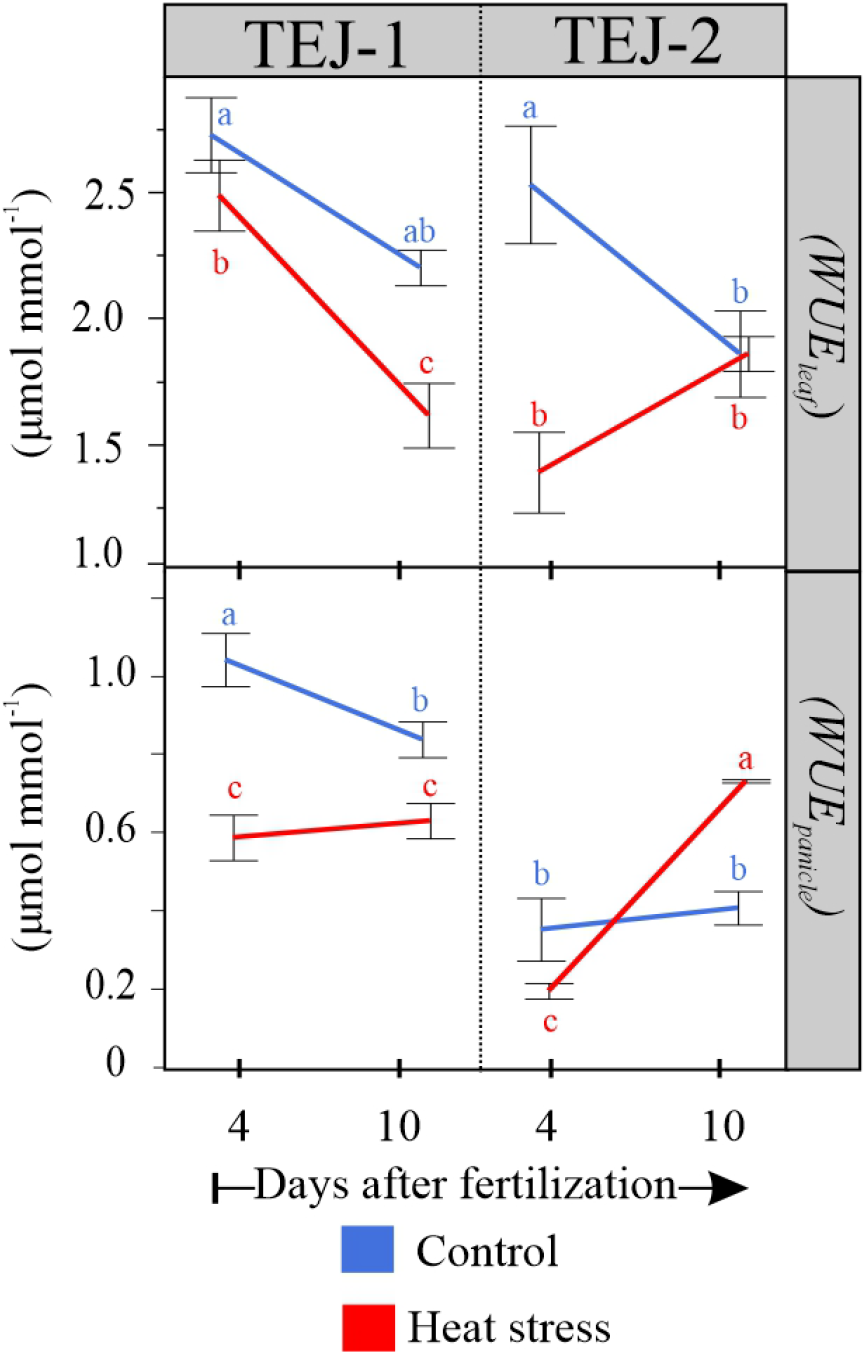
Water use efficiency measurements for leaf (*WUE*_*leaf*_) and panicle (*WUE*_*panicle*_) under control and HS for TEJ-1 and TEJ-2.

